# Cochlear dysfunction as an early biomarker for cognitive decline in normal hearing and mild hearing loss

**DOI:** 10.1101/2023.02.03.527051

**Authors:** Vicente Medel, Paul H. Delano, Chama Belkhiria, Alexis Leiva, Cristina De Gatica, Victor Vidal, Carlos F. Navarro, Simon San Martín, Melissa Martínez, Christine Gierke, Ximena García, Mauricio Cerda, Rodrigo Vergara, Carolina Delgado, Gonzalo Farías

## Abstract

Age-related hearing loss (presbycusis) at moderate levels (>40 dB HL) has been recognized as an important risk factor for cognitive decline. However, whether individuals with mild hearing loss (audiogram thresholds between 25 and 40 dB HL) or even those with normal audiograms (<25 dB HL) have a higher risk of dementia, is still debated. Importantly, these early stages of presbycusis are the most common among the elderly, indicating the need to screen and identify individuals with early presbycusis that have an increased risk of cognitive decline. Unfortunately, in this group of patients, audiogram thresholds are not sufficiently sensitive to detect all the hearing impairments that are related to cognitive decline. Consequently, at the individual level, audiogram thresholds are not good estimators of the dementia risk in the group with mild hearing loss or normal hearing thresholds. Here, we propose to use distortion product otoacoustic emissions (DPOAE), as an objective and sensitive tool to estimate the risk of clinically relevant cognitive decline in elders with normal hearing o mild hearing loss. We assessed neuropsychological, brain magnetic resonance imaging, and auditory analyses on 94 subjects aged >64 years old. In addition, cognitive and functional performance was evaluated with the Clinical Dementia Rating Sum of Boxes (CDR SoB), assessed through structured interviews conducted by neurologists, who were blind to the DPOAE results. We found that cochlear dysfunction, measured by DPOAE -and not by conventional audiometry-, was associated with CDR SoB classification and brain atrophy in the group with mild hearing loss (25 to 40 dB), and normal hearing (<25 dB). Our findings suggest that DPOAE may be a non-invasive tool for detecting neurodegeneration and cognitive decline in the elderly, potentially allowing for early intervention.

## Introduction

The prevalence of dementia is rapidly increasing, from 50 millions in 2015, to 150 millions in 2022, impacting global health systems worldwide (Alzheimer’s International Report 2020). A key challenge is the identification of reliable biomarkers for early diagnosis that could allow the implementation of prevention strategies for cognitive decline. In the last five years, hearing loss has emerged as one of the most relevant modifiable risk factors for dementia (1–4). The World Health Organization defines hearing loss as a 25 dB elevation in hearing thresholds, which are usually measured using perceptual audiometry, estimating a global prevalence of around 500 million people (WHO,2021). Evidence shows that the risk for cognitive decline and all-cause dementia increases with moderate hearing loss (greater than 40 dB HL) (5, 6), however, the relationship between hearing loss and cognitive decline could start earlier, including mild hearing loss (in the range between 25 and 40 dB), or even subjects with normal hearing thresholds (<25 dB HL)(7–9). Importantly, individuals with mild hearing loss (25-40 dB HL) or normal hearing thresholds (<25 dB HL) can have additional hearing impairments that are not detected by conventional audiometry, such as cochlear dead regions, cochlear synaptopathy or hidden hearing loss and central auditory processing disorder (10–12).

Hearing impairments can be estimated with subjective methods, such as audiometer tests and psychoacoustical tasks, and with objective methods, such as otoacoustic emissions or auditory evoked potentials (13). Otoacoustic emissions are inaudible low-level sounds that are emitted by the outer hair cells (OHC) of the cochlea (14). Its presence reflects normal cochlear functioning and OHC survival (15). Recently, we measured a subtype of otoacoustic emissions elicited by two tones, known as distortion product otoacoustic emissions (DPOAE, see Figure 1 and Methods section for more details on DPOAE measurements) and brain structural magnetic resonance imaging (MRI) in a group of aged subjects with mild hearing loss, evidencing significant associations between the loss of DPOAE and atrophy of non-auditory brain regions, including the cingulate cortex, insula and amygdala (16, 17). The structural alterations in the thickness and volume of cortical, and the volume of subcortical brain regions were related to cognitive and behavioral impairments in different domains, including face recognition and the executive function (16–18). However, whether these cognitive and behavioral impairments are related to the clinical phenotype of cognitive decline and dementia is unknown.

**Fig. 1.**
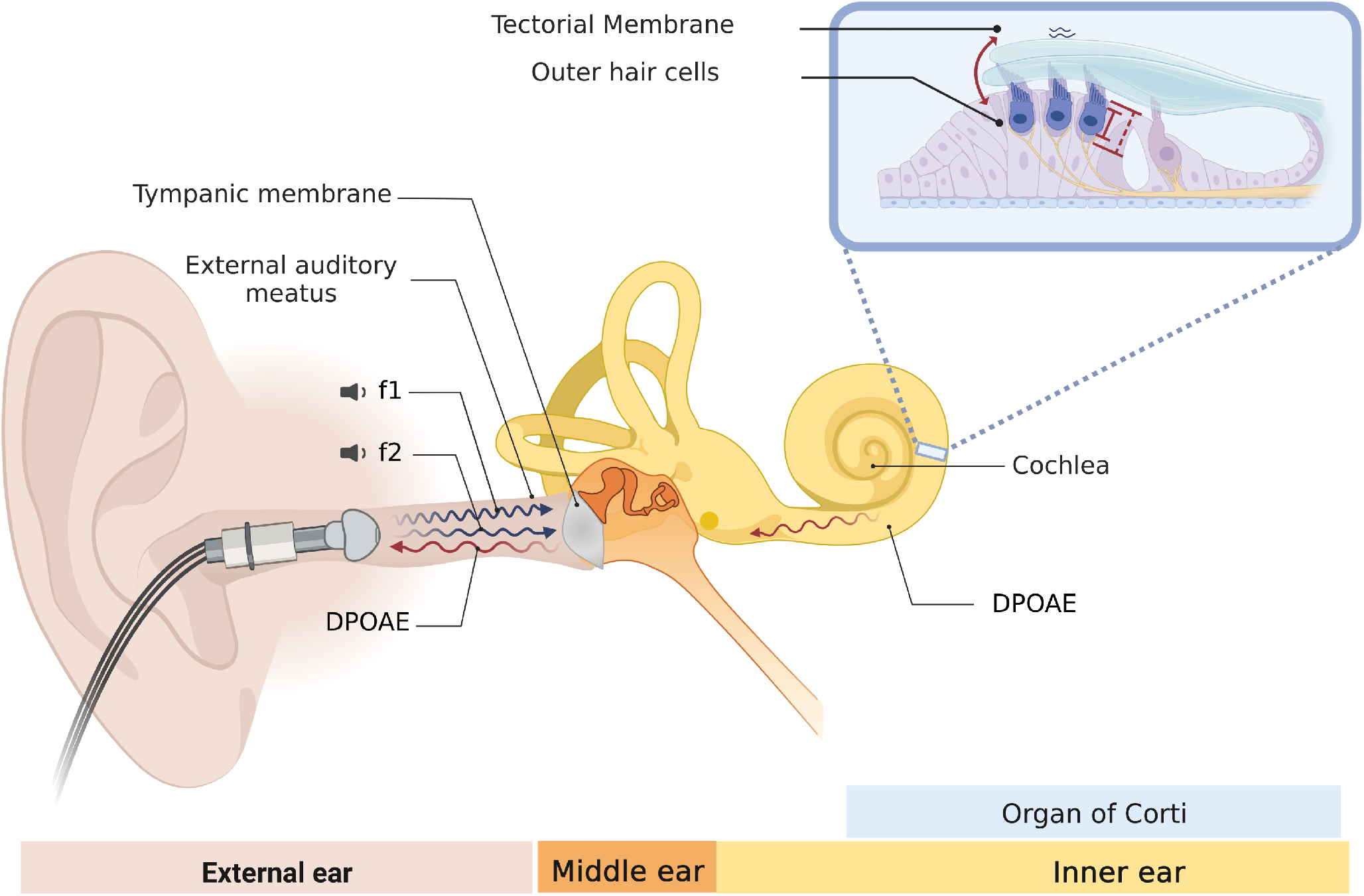
Distortion product otoacoustic emissions (DPOAE) are sounds emitted by normal cochlear hair cells. This figure describes the biological origin of DPOAEs, and the methods used to measure them. A microphone probe with two speakers is inserted and sealed in the external ear canal. Two tones of different frequencies (f1 and f2) are presented independently through the two speakers. These two tones are transmitted across the external and middle ear, reaching the cochlear receptor. In the inner ear (cochlear receptor), the two tones generate mechanical distortions at different positions of the basilar membrane of the cochlea, including the 2f1-f2 position, which is the most widely used DPOAE on clinical and research settings. The presence of DPOAE depends on the normal functioning of outer hair cells (OHC). OHCs possess electromotily, a physiological mechanism in which these cells transduce membrane voltage changes into mechanical vibrations, a physiological process known as the “cochlear amplifier”. As a result of this biological amplification, f1 and f2 tones interact and generate mechanical distortions at different cochlear positions, including the 2f1-f2 position. These distortions travel back to the external ear where they can be recorded with a sensitive microphone and measured as a DPOAE at a certain frequency and amplitude in dB SPL.

Here, we hypothesized that the loss of DPOAE is associated with the clinical phenotype of cognitive decline. To test our hypothesis, we studied the presence of DPOAE as a proxy of cochlear functioning and OHC survival (as in Belkhiria et al. (16)), while the cognitive clinical profile was independently assessed by blind experienced neurologists determining the Clinical Dementia Rating Sum of Boxes (CDR SoB; (19)). In addition, subjects were evaluated with comprehensive audiological, neuropsychological, and brain MRI evaluations.

## Results

A total of 94 subjects (65 female) with an average age of 72.7 ± 5 years (mean, SD), and mean education level of 9.5 ±5.1 years were obtained from the ANDES cohort (16, 17). Data from this cohort includes DPOAE measurements, audiogram thresholds, assessed by Pure-Tone Average (PTA), a battery of neuropsychological tests for cognitive decline, and structural brain MRI at 3-Tesla.

The mean PTA of the 94 individuals was 24.3 ± 8 dB HL, including 32 subjects with normal hearing thresholds (<25 dB HL) and 62 with mild hearing loss (between 25 and 40 dB HL). None of the individuals used hearing aids at the moment of evaluations. Regarding DPOAE, we calculated the total number of detected DPOAE in eight different frequencies in both ears (range between 0 and 16, bigger is better, see methods section), yielding an average number of 8.4 ± 5 detected DPOAEs per individual.

As DPOAEs have been used as a proxy of hearing sensitivity (Shaffer et al., 2003), we explored the level of correlation between DPOAE and PTA variables in our data. DPOAE measurements and PTA audiogram thresholds partly converge (Spearman’s rank correlation *ρ* = −0.64, p-value < 0.0001; Figure 2A), suggesting that they capture similar but not equal auditory characteristics. In relation to cognitive and neurological variables, the mean Mini-Mental State Examination (MMSE) score was 27 ± 3.4, while the mean CDR SoB was 0.94 ± 2.3. A summary of demographic data is reported in Table 1.

**Fig. 2.**
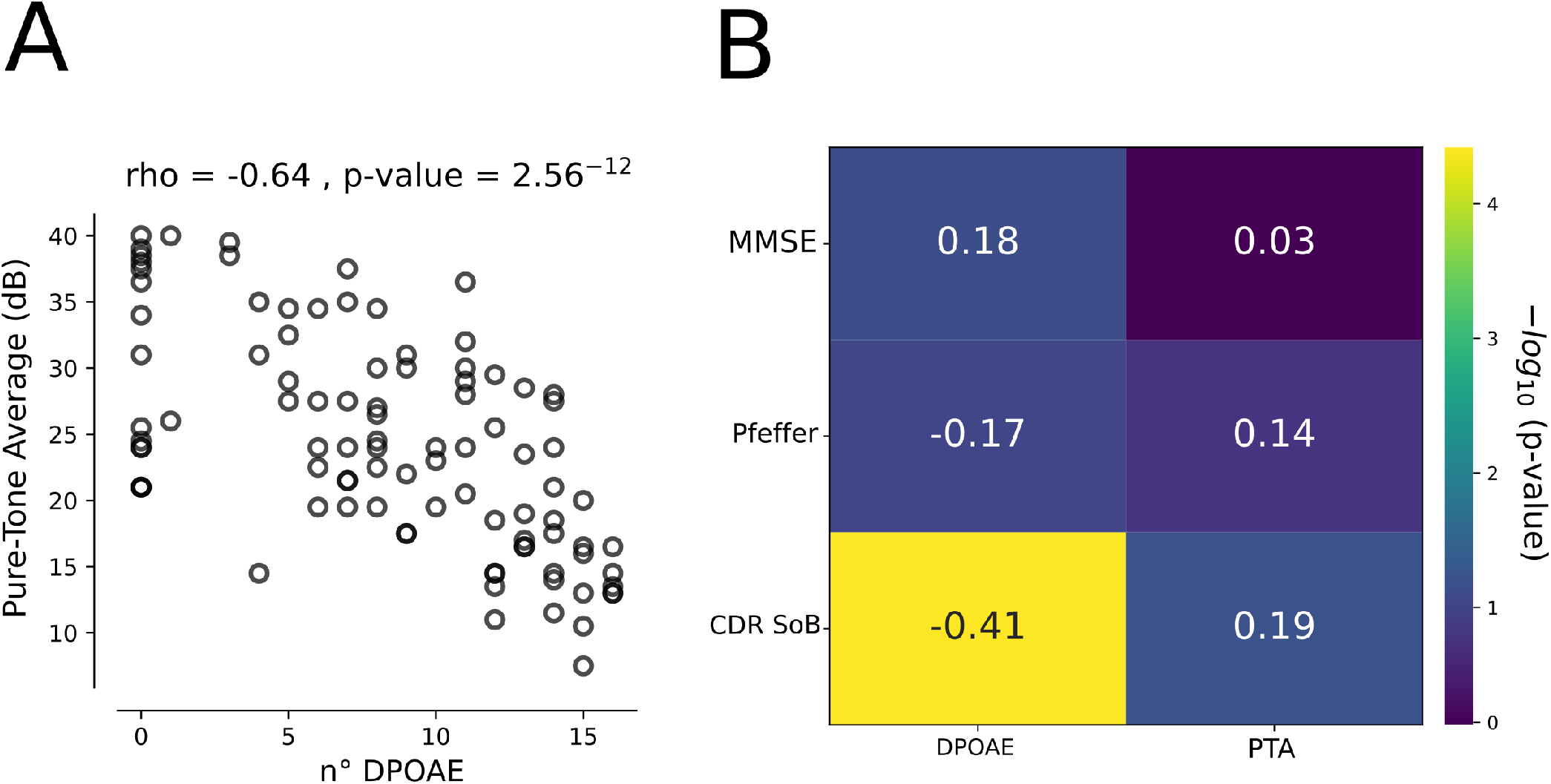
Differential contributions DPOAE and PTA over cognition. (**A**) Pure-Tone Average (PTA) negatively correlates with the number of DPOAE, evidencing that a larger number of DPOAE is correlated with better PTA (**B**) Correlation matrix with annotated Spearman’s rho values in each cell, between audiological measures and Mini-mental (MMSE), the Pfeffer questionnaire, and the CDR SoB. Note that only the correlation between CDR SoB and the number of DPOAE was significant (p<0.001). Color-bar in the y-axis represents the −*log*_10_ of the p-value obtained from Spearman’s Rank correlation.

**Table 1.**
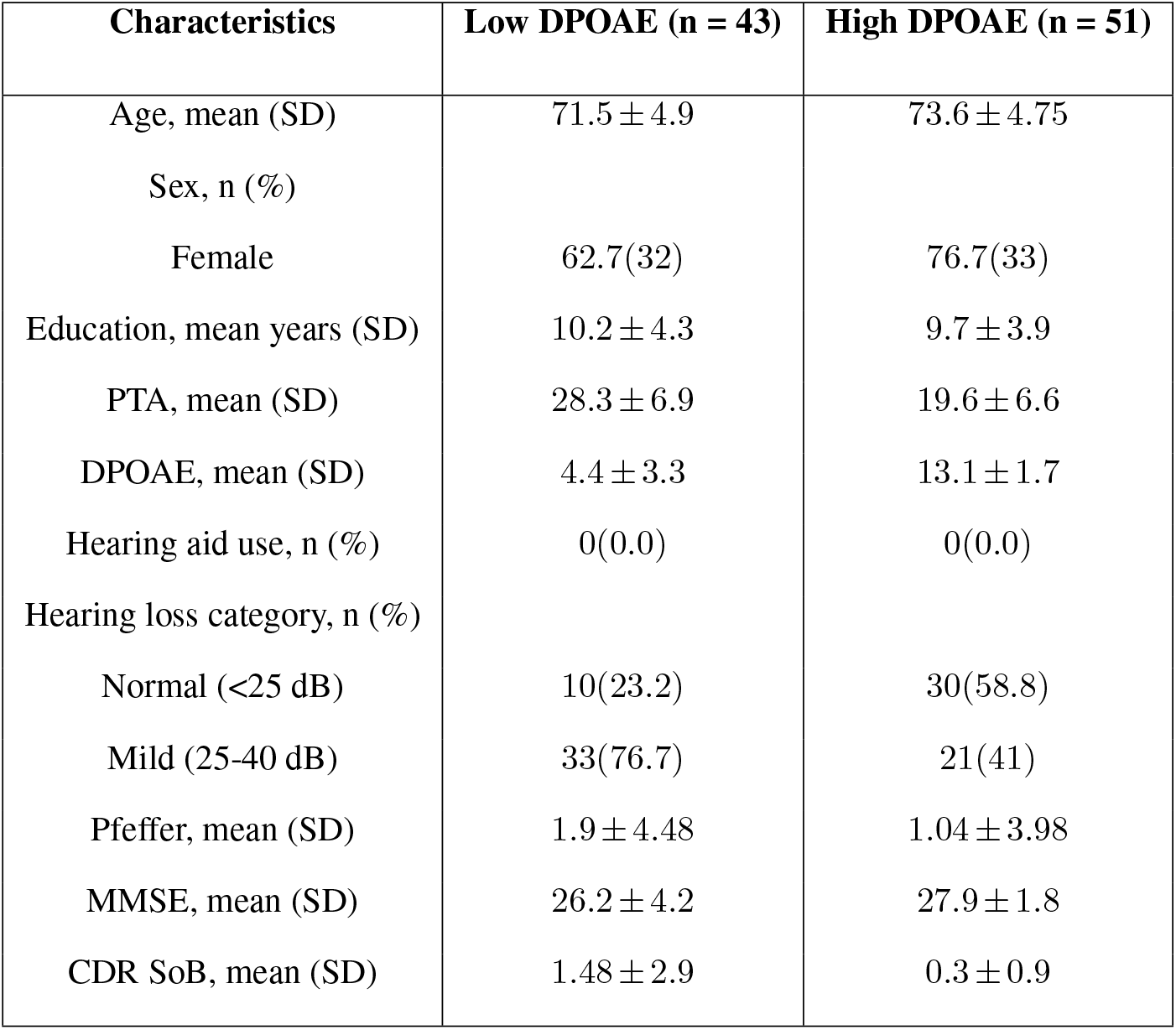
Audiologial and Neurological profile of the studied subjects (n=94)

| Characteristics | Low DPOAE (n = 43) | High DPOAE (n = 51) |
| --- | --- | --- |
| Age, mean (SD) | 71.5 ± 4.9 | 73.6 ± 4.75 |
| Sex, n (%) |  |  |
| Female | 62.7(32) | 76.7(33) |
| Education, mean years (SD) | 10.2 ± 4.3 | 9.7 ± 3.9 |
| PTA, mean (SD) | 28.3 ± 6.9 | 19.6 ± 6.6 |
| DPOAE, mean (SD) | 4.4 ± 3.3 | 13.1 ± 1.7 |
| Hearing aid use, n (%) | 0(0.0) | 0(0.0) |
| Hearing loss category, n (%) |  |  |
| Normal (<25 dB) | 10(23.2) | 30(58.8) |
| Mild (25-40 dB) | 33(76.7) | 21(41) |
| Pfeffer, mean (SD) | 1.9 ± 4.48 | 1.04 ± 3.98 |
| MMSE, mean (SD) | 26.2 ± 4.2 | 27.9 ± 1.8 |
| CDR SoB, mean (SD) | 1.48 ± 2.9 | 0.3 ± 0.9 |

### Less DPOAE is correlated with worse dementia rating

Next, we analyzed the possible relations between the number of DPOAE and PTA with cognitive traits, such as global cognitive performance evaluated through MMSE, functional status quantified using Pfeffer, and the CDR SoB, which was obtained by clinical neurological evaluations that were blind to the DPOAE results, providing a quantitative index of the level of cognitive impairment as a dementia rating. Neither MMSE nor Pfeffer presented a significant correlation with the number of DPOAEs (MMSE Spearman’s Rank Correlation, *ρ* = 0.18, p-value = 0.075 and Pfeffer Spearman’s Rank Correlation, *ρ* = −0.17, p-value = 0.09) and PTA (MMSE Spearman’s rank correlation, *ρ* = 0.03, p-value = 0.74 and Pfeffer Spearman’s Rank Correlation, *ρ* = 0.14, p-value = 0.16). Interestingly, CDR SoB showed a significant correlation with the number of DPOAE (Spearman’s Rank Correlation *ρ* = −0.41, p-value < 0.00001) while there was a non-significant correlation between PTA and CDR SoB score (Spearman’s Rank Correlation *ρ* = 0.19, p-value = 0.06). These results suggest that, although the number of DPOAE and PTA have a partial statistical convergence (Figure 2A), they correlated diferentially with dementia rating, suggesting that DPOAE is a more sensitive hearing assessment for evidencing the relation between hearing impairments and clinical profile of cognitive decline related to the CDR SoB score (Figure 2B).

### Neuroimaging biomarkers correlate with DPOAE

Currently, there are well-validated brain MRI biomarkers that are commonly used for the clinical evaluation of patients with cognitive decline complaints, such as the volumetric changes of the hippocampus and lateral ventricles. To test if cochlear dysfunction is associated to these cognitive decline neuroimaging biomarkers, we first divided subjects by their number of DPOAE, calculating the median value of the whole population, thus defining low DPOAE and high DPOAE groups. Strikingly, we found that the volume of bilateral hippocampus was significantly more atrophied in the low DPOAE group as compared to the high DPOAE group (Figure 3A, left panel, Mann-Whitney U test p-value = 0.0015). In addition, we found that bilateral lateral ventricles were significantly larger in the low DPOAE group (Figure 3B, left panel. Mann-Whitney U test p-value = 0.00003). There was a significant correlation between the number of DPOAE with the volume of both hippocampus (Spearman’s Rank Correlation, *ρ* = 0.312, p = 0.002, Figure 3A, right panel) and lateral ventricles (Spearman’s Rank Correlation *ρ* = −0.37, p = 0.0001, Figure 3B, right panel).

**Fig. 3.**
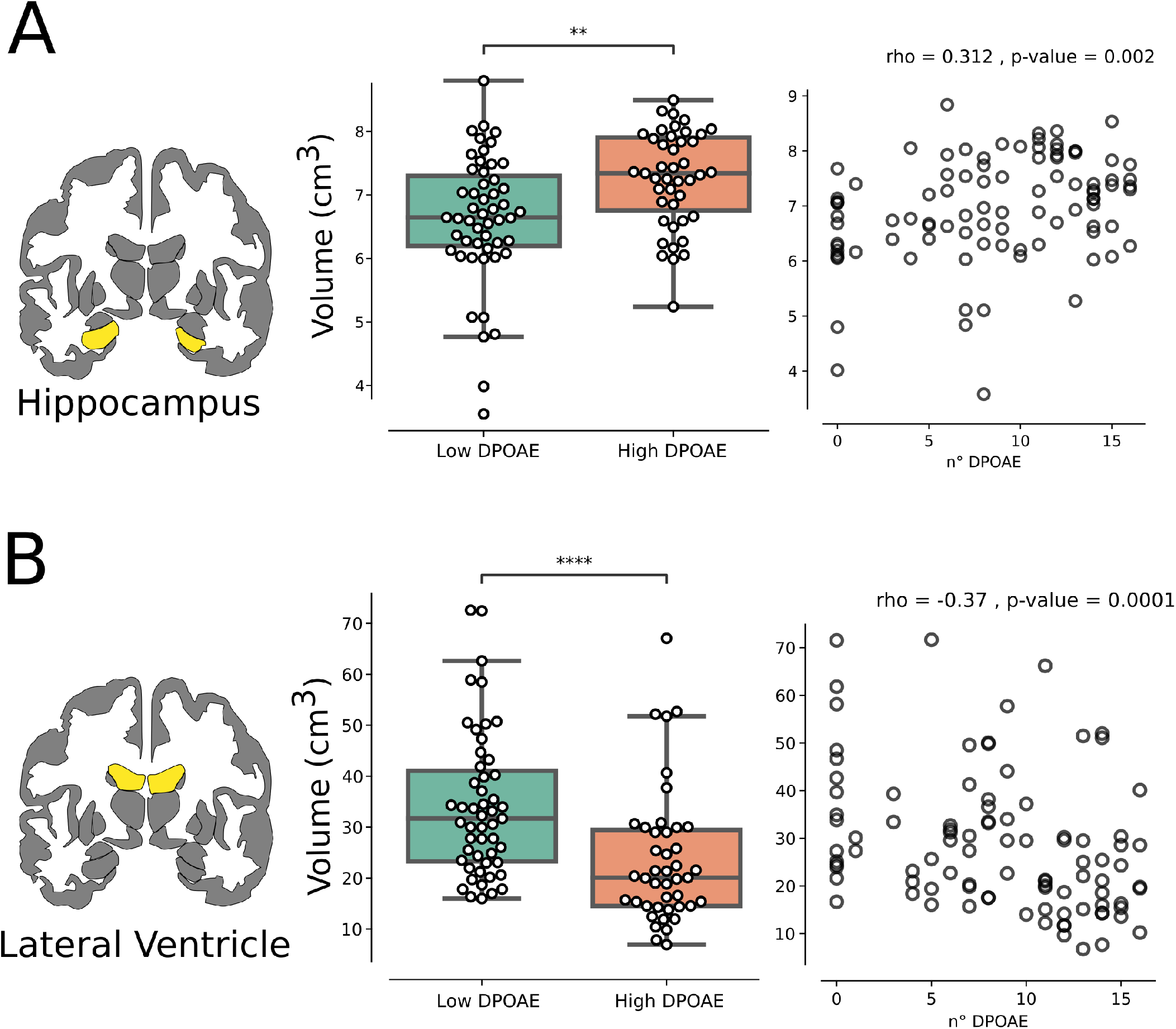
DPOAE correlates with neuroimaging biomarkers of cognitive decline. (**A**) In left, a coronal view of bilateral Hippocampal volume. In middle, total hippocampal volume is more atrophied in Low DPOAE group as compared to high DPOAE group (Mann-Whitney U test p-value = 0.001). In right, a significant correlation between total hippocampal volume and the number of DPOAE shows a significant positive correlation. (**B**) In left, a coronal view of bilateral Lateral Ventricle volume. In middle, total ventricular volume is more hypertrophied in Low DPOAE group as compared to high DPOAE group (p-value = 0.00003). In right, a significant correlation between total ventricular volume and the number of DPOAE shows a significant negative correlation. All correlations are Spearman’s rank.

### DPOAE predicts cognitive decline

To test if cochlear dysfunction as measured by DPOAE predicts cognitive decline in our cohort and if overcomes conventional audiometry as measured by PTA, we evaluated its capacity to discriminate between control (CDR SoB < 0.5) and risk of cognitive decline (CDR SoB >= 0.5). Receiver operating characteristic (ROC) curve analysis was performed and the area under the curve (AUC) was calculated. Fivefold cross-validation was performed for PTA (AUC = 0.45 ± 0.26, Figure 4A) showing its prediction power is near random. In contrast, the number of DPOAE predicted with good discriminability (AUC = 0.81 ± 0.1, Figure 4A). To evaluate if DPOAE consistently predicts better than PTA, we obtained 1000 bootstraped AUC score for each term, and statistically compared the scores. Notably, PTA significantly predicts worst than DPOAE (Mann-Whitney U test p-value < 0.0001), and DPOAE shows low AUC variability. These results show that DPOAE overcomes PTA in terms of discriminability (Figure 4B), and robustly predicts the risk of clinically relevant cognitive decline in a cohort of normal and mild hearing loss elders.

**Fig. 4.**
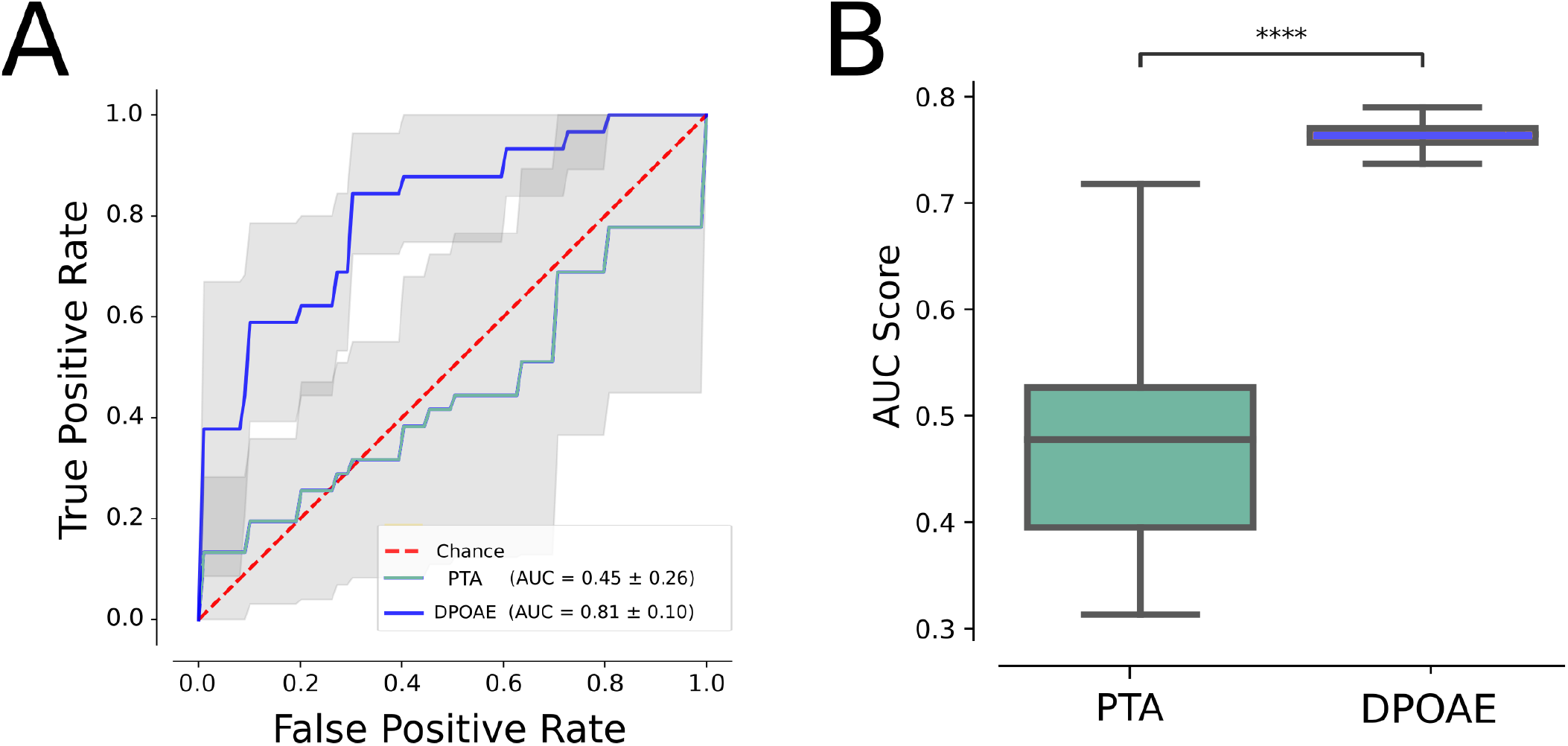
DPOAE and not PTA predicts cognitive impairment in normal and mild hearing loss individuals. (**A**) Five-fold cross-validation using PTA (green, AUC=0.45) and DPOAE (blue, AUC=0.81) to predict cognitive impairment (CDR SoB > 0.5). (**B**) Boxplot comparing 1000 bootstrap AUC scores, depicting that PTA values significantly predict worst than DPOAE (Mann–Whitney U test p-value < 0.0001)

## Discussion

We found that in a sample of normal hearing (PTA < 25 dB) and age-related mild-hearing loss individuals (PTA between 25 and 40 dB HL), the number of DPOAE is significantly correlated with the clinical classification of dementia (CDR SoB scale) evaluated by expert neurologists that were blind to the DPOAE results. In the same group of subjects, audiometric hearing thresholds were not correlated with the CDR SoB scale. These findings suggest that evaluating cochlear OHC function by means of DPOAE detection is a more sensitive test than audiogram thresholds to estimate clinical-relevant cognitive impairment risk in elders. Importantly, these results stress the fact that, although DPOAE and PTA are significantly correlated (Figure 2A), they possess relevant differences. For instance, audiometric PTA is obtained by subjective behavioral responses to pure tones, which can be influenced by multiple sources of variability, such as attention and the cognitive load level(20), while DPOAE is an objective measure that does not need subject cooperation and reflects cochlear integrity. In addition, and concordant with our previous works (16, 17), we found significant correlations between the loss of DPOAE and the volume of different brains structures, such as bilateral hippocampus and lateral ventricle volumes (Figure 3).

### What are the mechanisms that relate DPOAE loss with brain atrophy and risk of cognitive decline?

There are at least three possible mechanisms relating DPOAE loss to the risk of dementia. First, DPOAE is a sensitive and objetive measure to detect hearing impairment and is highly correlated with PTA thresholds. Thus, DPOAE could be considered as a more sensitive measure to estimate hearing impairments than audiometric PTA (Figure 4). Second, as a proxy of OHC loss, DPOAE loss might reflect the process of cellular aging due to non-specific neurodegenerative and vascular damage of cells located inside the cranium, and in an speculative statement, DPOAE loss might be a general estimator of neuronal survival in the brain. Finally, DPOAE presence can also be affected by acoustic trauma (21), and there is evidence in animal models that acoustic trauma can induce hippocampal atrophy which is also related to neurodegenerative tau pathology and β-amyloid in the brain (22, 23). Therefore, different from PTA thresholds, DPOAE loss might be reflecting a number of biological processes that can contribute in additive ways to cognitive decline in elders, which could explain the better performance in the AUC score for classifying CDR SoB scores in this cohort of normal hearing and mild hearing loss individuals (Figure 4).

It is noteworthy that DPOAE is related to cognitive impairment evaluated by CDR SoB, a scale that measures various cognitive and functional domains but does not show a significative relationship with MMSE, which measures memory and cognition, or with Pfeffer, which evaluates functional activities. MMSE has a low sensibility to predict the presence of initial or mild cognitive impairment and has a better performance in dementia cases Our findings may have two practical implications for future research. First, from a pathophysiological point of view, DPOAE loss as a measure of cochlear OHC damage, reinforces the connection between degeneration of auditory structures with broader neurodegenerative phenomena such as those seen in Alzheimer’s disease and related dementias. Secondly, from the clinical point of view, it highlights the potential of DPOAE as a possible biomarker of neurodegenerative phenomena and cognitive impairment. The determination of DPOAE can constitute an objective, simple and accessible biomarker to detect elderly people at risk of cognitive deterioration, considering the AUC over 0.8 in Figure 4. This idea is supported by recent findings in animal models showing a significant relation between auditory functions and tau protein levels in the cerebrospinal fluid (24).

The practical implementation of using DPOAE as an screening tool for detecting elders at higher risk of developing cognitive decline is supported by the routinely use of DPOAE in newborn hearing screening programs worldwide (25). Equipment for DPOAE measurements is available in hospitals in several countries, including universal screening programs (26–28). Audiologists or other professionals are already trained in the measurement of DPOAE, and the examination is relatively simple, fast, and available at a reasonable cost. There are experiences of DPOAE use for the screening of hearing loss in adults and of its good correlation with other audiological measurements such as brainstem evoked auditory responses (29). From our work, we propose that DPOAE detection can be used in elders, for estimating the risk of cognitive decline at an early stage, including the group of subjects with normal hearing or mild hearing loss.

## Materials and Methods

### Subjects

A total of 94 older adults from the Auditory and Dementia study (ANDES) cohort (16) were recruited according to the inclusion and exclusion criteria for this study. The patients were recruited from Recoleta’s primary health public center in Santiago de Chile. All of the included subjects gave written informed consent in accordance with the Declaration of Helsinki. The inclusion and exclusion criteria were the following: (i) age over 65 years at the time of recruitment, (ii) no history of neurological or psychiatric illness, (iii) no causes of hearing loss different from presbycusis (e.g., conductive hearing loss), (iv) no patients using hearing aids. All of the undertaken actions were approved by the Ethics Committee of the Clinical Hospital of the University of Chile with permission number OAIC752/15.

### Clinical Assessment

Clinical assessment included: comprehensive cognitive assessment including the Mini-Mental State Examination (MMSE) (30), Functional status as quantified by Pfeffer’s Functional Activities Questionnaire (31), and Clinical Dementia Rating Sum of Boxes for assessing the severity of cognitive and functional impairments associated with dementia (19, 32). Routine neurological and psychiatric examination included daily living questionnaire filled in by close relative or partner interviewed by a neuropsychologist.

### Audiological Evaluations

Evaluations were carried out in the Otolaryngology Department of the Clinical Hospital of the University of Chile. Air conduction pure tone audiometric hearing thresholds were evaluated at 0.125, 0.25, 0.5, 1, 2, 3, 4, 6, and 8 kHz for each subject in both ears using a clinical audiometer (AC40, Interacoustics®). Bone conduction thresholds were measured at 0.25, 0.5, 1, 2, 3 and 4 kHz to rule out conductive hearing loss. Pure Tone Average (PTA) at 0.5, 1, 2, and 4 kHz was calculated for each subject in both ears. Subjects were classified according to their hearing level: normal hearing (≤ 25dB), mild presbycusis (*>* 25 dB, and ≤ 40 dB) based on the average PTA score of both ears.

### Distortion product otoacoustic emissions (DPOAE)

DPOAE was evaluated as described by (16). Briefly, DPOAE (2f1–f2) was measured as a proxy of the cochlear amplifier function, using an ER10C microphone (Etymotic Research®), presenting eight pairs of primary tones (f1 and f2, at 65 and 55 dB SPL, f2/f1 ratio of 1.22) in each ear at eight different 2f1–f2 frequencies: 707 Hz, 891 Hz, 1122 Hz, 1414 Hz, 1781 Hz, 2244 Hz, 2828 Hz, and 3563 Hz. Importantly, using different pairs of tones (f1 and f2) at different frequencies, allows the measurement of the 2f1-f2 DPOAE at different cochlear positions that can be used as a proxy of hearing impairments and cochlear hair cell survival in aged subjects. The amplitude of a DPOAE (dB SPL) should be at least 6 dB above the noise floor. The number of detectable DPOAEs per ear -i.e. 0 to 8-was counted and a number of “0” -no detectable DPOAE in that ear- was considered as cochlear amplifier dysfunction while “8” implied normal cochlear amplifier function. We used the total number of detected DPOAE of both ears (range from 0 to 16).

### Image acquisition

Imaging data were acquired using a MAGNETOM Skyra 3-Tesla whole-body MRI Scanner (Siemens Healthcare GmbHR, Erlangen, Germany) using a T1-MPRAGE sequence. Contiguous images across the entire brain were acquired with the following parameters: time echo (TE) = 232 ms, time repetition (TR) = 2300 ms, flip angle = 8, 26 slices, matrix = 256×256, voxel size = 0.94×0.94×0.9 mm3. We also registered T2-weighted turbo spin echo (TSE) (4500 TR ms, 92 TE ms) and fluid-attenuated inversion recovery (FLAIR) (8000 TR ms, 94 TE ms, 2500 TI ms) to inspect structural abnormalities. The acquisition duration was 30 min with a total of 440 images for each subject.

### Image preprocessing and analysis

The morphometric analysis was carried out by FreeSurfer, version 6 running under Centos 6. A single Linux workstation was used for the T1-weighted image analysis of individual subjects as suggested by (33). The FreeSurfer processes cortical reconstruction (34) through several steps: volume registration with the Talairach atlas, bias field correction, initial volumetric labeling, non-linear alignment to the Talairach space, and final volume labeling. Briefly, the automatic “recon-all” function produces representations of the cortical surfaces. It uses both intensity and continuity information from the entire threedimensional MR volume in segmentation and deformation procedures. It creates gross brain volume extents for larger-scale regions (i.e., the total number of voxels per region): total gray and white matter, subcortical gray matter, brain mask volume, and estimated total intracranial volume. The reliability between manual tracing and automatic volume measurements has been validated. The accordance between manual tracings and automatically obtained segmentations was similar to the agreement between manual tracings (35). All volumes were visually inspected, and if needed, edited by a trained researcher according to standard processes.

We selected regions of interest that have been consistently implicated in previous neuroimaging studies relating to audition, cognition, and dementia, such as Hippocampus and the Lateral Ventricles.

### Statistical Analysis

Spearman’s Rank where used to asses the correlation between variables. For brain volume comparison between High and Low DPOAE groups a Mann-Whitney U test was used. ROC curves and AUC scores were obtained using scikit-learn package (36). For the comparison of PTA and DPOAE scores, a 1000 bootstrapping approach was used, and a Mann-Whitney U test for statistical comparison.

## ACKNOWLEDGEMENTS

This research work was supported by the National Agency for Research and Development of Chile (ANID), FONDEF ID20I10371, FONDECYT 1221696, FONDECYT 1220607, FONDEQUIP EQM210020, Fondo Basal ANID FB0008 to PHD, Centro Nacional de Inteligencia Artificial CENIA, FB210017, BASAL, ANID to RV, Proyecto Milenio ICN09_015, and Fundación Guillermo Puelma.

## Bibliography

1. Gill Livingston, Andrew Sommerlad, Vasiliki Orgeta, Sergi G Costafreda, Jonathan Huntley, David Ames, Clive Ballard, Sube Banerjee, Alistair Burns, Jiska Cohen-Mansfield, et al. Dementia prevention, intervention, and care. The Lancet, 390(10113):2673–2734, 2017.

2. Gill Livingston, Jonathan Huntley, Andrew Sommerlad, David Ames, Clive Ballard, Sube Banerjee, Carol Brayne, Alistair Burns, Jiska Cohen-Mansfield, Claudia Cooper, et al. Dementia prevention, intervention, and care: 2020 report of the lancet commission. The Lancet, 396(10248):413–446, 2020.

3. Katharine K Brewster, Jennifer A Deal, Frank R Lin, and Bret R Rutherford. Considering hearing loss as a modifiable risk factor for dementia. Expert review of neurotherapeutics, 22(9):805–813, 2022.

4. Rodrigo C Vergara, Pedro Zitko, Andrea Slachevsky, Consuelo San Martin, and Carolina Delgado. Population attributable fraction of modifiable risk factors for dementia in chile. Alzheimer’s & Dementia: Diagnosis, Assessment & Disease Monitoring, 14(1):e12273, 2022.

5. Jennifer A Deal, Josh Betz, Kristine Yaffe, Tamara Harris, Elizabeth Purchase-Helzner, Suzanne Satterfield, Sheila Pratt, Nandini Govil, Eleanor M Simonsick, Frank R Lin, et al. Hearing impairment and incident dementia and cognitive decline in older adults: the health abc study. Journals of Gerontology Series A: Biomedical Sciences and Medical Sciences, 72(5):703–709, 2017.

6. Natalia Tamblay, Dorothy Boggs, Barbara Huidobro, Daniel Tapia-Mora, Katherine Anabalon, Carolina Delgado, Sarah Polack, Tess Bright, and Mariela C Torrente. Prevalence of cognitive impairment and its association with hearing loss among adults over 50 years of age: Results from a population-based survey in santiago, chile. American Journal of Audiology, pages 1–10, 2023.

7. Jeremy CS Johnson, Charles R Marshall, Rimona S Weil, Doris-Eva Bamiou, Chris JD Hardy, and Jason D Warren. Hearing and dementia: from ears to brain. Brain, 144(2): 391–401, 2021.

8. Alexandria L Irace, Nicole M Armstrong, Jennifer A Deal, Alexander Chern, Luigi Ferrucci, Frank R Lin, Susan M Resnick, and Justin S Golub. Longitudinal associations of subclinical hearing loss with cognitive decline. The Journals of Gerontology: Series A, 77(3):623–631, 2022.

9. Alexander Chern, Alexandria L Irace, Rahul K Sharma, Yuan Zhang, Qixuan Chen, and Justin S Golub. The longitudinal association of subclinical hearing loss with cognition in the health, aging and body composition study. Frontiers in aging neuroscience, 13, 2021.

10. Rodolfo Sardone, Petronilla Battista, Rossella Donghia, Madia Lozupone, Rosanna Tortelli, Vito Guerra, Alessandra Grasso, Chiara Griseta, Fabio Castellana, Roberta Zupo, et al. Age-related central auditory processing disorder, mci, and dementia in an older population of southern italy. Otolaryngology–head and Neck Surgery, 163(2):348–355, 2020.

11. Vinay, Sandhya, and Brian CJ Moore. Effect of age, test frequency and level on thresholds for the ten (hl) test for people with normal hearing. International Journal of Audiology, 59 (12):915–920, 2020.

12. Munir Demir Bajin, Valerie Dahm, and Vincent YW Lin. Hidden hearing loss: current concepts. Current Opinion in Otolaryngology & Head and Neck Surgery, 30(5):321–325, 2022.

13. RA Davies. Audiometry and other hearing tests. Handbook of clinical neurology, 137:157–176, 2016.

14. David T Kemp. Stimulated acoustic emissions from within the human auditory system. The Journal of the Acoustical Society of America, 64(5):1386–1391, 1978.

15. Lauren A Shaffer, Robert H Withnell, Sumit Dhar, David J Lilly, Shawn S Goodman, and Kelley M Harmon. Sources and mechanisms of dpoae generation: implications for the prediction of auditory sensitivity. Ear and hearing, 24(5):367–379, 2003.

16. Chama Belkhiria, Rodrigo C Vergara, Simón San Martín, Alexis Leiva, Bruno Marcenaro, Melissa Martinez, Carolina Delgado, and Paul H Delano. Cingulate cortex atrophy is associated with hearing loss in presbycusis with cochlear amplifier dysfunction. Frontiers in aging neuroscience, 11:97, 2019.

17. Chama Belkhiria, Rodrigo C Vergara, Simón San Martin, Alexis Leiva, Melissa Martinez, Bruno Marcenaro, Maricarmen Andrade, Paul H Delano, and Carolina Delgado. Insula and amygdala atrophy are associated with functional impairment in subjects with presbycusis. Frontiers in aging neuroscience, 12:102, 2020.

18. Chama Belkhiria, Rodrigo C Vergara, Melissa Martinez, Paul H Delano, and Carolina Delgado. Neural links between facial emotion recognition and cognitive impairment in presbycusis. International Journal of Geriatric Psychiatry, 36(8):1171–1178, 2021.

19. Sid E O’Bryant, Stephen C Waring, C Munro Cullum, James Hall, Laura Lacritz, Paul J Massman, Philip J Lupo, Joan S Reisch, Rachelle Doody, Texas Alzheimer’s Research Consortium, et al. Staging dementia using clinical dementia rating scale sum of boxes scores: a texas alzheimer’s research consortium study. Archives of neurology, 65(8):1091–1095, 2008.

20. Antje Heinrich, Melanie A Ferguson, and Sven L Mattys. Effects of cognitive load on puretone audiometry thresholds in younger and older adults. Ear and hearing, 41(4):907, 2020.

21. Roger P Hamernik, William A Ahroon, Brenda M Jock, and Jessie A Bennett. Noise-induced threshold shift dynamics measured with distortion-product otoacoustic emissions and auditory evoked potentials in chinchillas with inner hair cell deficient cochleas. Hearing research, 118(1-2):73–82, 1998.

22. Fabiola Paciello, Marco Rinaudo, Valentina Longo, Sara Cocco, Giulia Conforto, Anna Pisani, Maria Vittoria Podda, Anna Rita Fetoni, Gaetano Paludetti, and Claudio Grassi. Auditory sensory deprivation induced by noise exposure exacerbates cognitive decline in a mouse model of alzheimer’s disease. Elife, 10, 2021.

23. Mengmeng Zheng, Jiangyu Yan, Wenjuan Hao, Yuan Ren, Ming Zhou, Yunzhi Wang, and Kai Wang. Worsening hearing was associated with higher *β*-amyloid and tau burden in age-related hearing loss. Scientific Reports, 12(1):1–7, 2022.

24. Wei Xu, Can Zhang, Jie-Qiong Li, Chen-Chen Tan, Xi-Peng Cao, Lan Tan, Jin-Tai Yu, et al. Age-related hearing loss accelerates cerebrospinal fluid tau levels and brain atrophy: a longitudinal study. Aging (Albany NY), 11(10):3156, 2019.

25. Thomas Janssen. A review of the effectiveness of otoacoustic emissions for evaluating hearing status after newborn screening. Otology & Neurotology, 34(6):1058–1063, 2013.

26. Heather L Porter, Stephen T Neely, and Michael P Gorga. Using benefit-cost ratio to select universal newborn hearing screening test criteria. Ear and hearing, 30(4):447, 2009.

27. Yojana Sharma, Girish Mishra, Sushen H Bhatt, and Somashekhar Nimbalkar. Neonatal hearing screening programme (nhsp): At a rural based tertiary care centre. Indian Journal of Otolaryngology and Head & Neck Surgery, 67(4):388–393, 2015.

28. Jacqueline K Bezuidenhout, Katijah Khoza-Shangase, Tim De Maayer, and Renate Strehlau. Outcomes of newborn hearing screening at an academic secondary level hospital in johannesburg, south africa. South African Journal of Communication Disorders, 68 (1):741, 2021.

29. Vidya Ramkumar, CS Vanaja, James W Hall, K Selvakumar, and Roopa Nagarajan. Validation of dpoae screening conducted by village health workers in a rural community with real-time click evoked tele-auditory brainstem response. International Journal of Audiology, 57(5):370–375, 2018.

30. Marshal F Folstein. A practical method for grading the cognitive state of patients for the clinician. J Psychiatr res, 12:189–198, 1992.

31. Robert I Pfeffer, Tom T Kurosaki, CH Harrah Jr, Jeffrey M Chance, and S Filos. Measurement of functional activities in older adults in the community. Journal of gerontology, 37(3): 323–329, 1982.

32. Charles P Hughes, Leonard Berg, Warren Danziger, Lawrence A Coben, and Ronald L Martin. A new clinical scale for the staging of dementia. The British journal of psychiatry, 140(6):566–572, 1982.

33. Ed HBM Gronenschild, Petra Habets, Heidi IL Jacobs, Ron Mengelers, Nico Rozendaal, Jim Van Os, and Machteld Marcelis. The effects of freesurfer version, workstation type, and macintosh operating system version on anatomical volume and cortical thickness measurements. PloS one, 7(6):e38234, 2012.

34. Bruce Fischl and Anders M Dale. Measuring the thickness of the human cerebral cortex from magnetic resonance images. Proceedings of the National Academy of Sciences, 97 (20):11050–11055, 2000.

35. Bruce Fischl, David H Salat, Evelina Busa, Marilyn Albert, Megan Dieterich, Christian Haselgrove, Andre Van Der Kouwe, Ron Killiany, David Kennedy, Shuna Klaveness, et al. Whole brain segmentation: automated labeling of neuroanatomical structures in the human brain. Neuron, 33(3):341–355, 2002.

36. F. Pedregosa, G. Varoquaux, A. Gramfort, V. Michel, B. Thirion, O. Grisel, M. Blondel, P. Prettenhofer, R. Weiss, V. Dubourg, J. Vanderplas, A. Passos, D. Cournapeau, M. Brucher, M. Perrot, and E. Duchesnay. Scikit-learn: Machine learning in Python. Journal of Machine Learning Research, 12:2825–2830, 2011.

